# Leveraging a physiologically based quantitative translational modeling platform for designing bispecific T cell engagers for treatment of multiple myeloma

**DOI:** 10.1101/2021.12.06.471352

**Authors:** Tomoki Yoneyama, Mi-Sook Kim, Konstantin Piatkov, Haiqing Wang, Andy Z.X. Zhu

**Author notes:** Corresponding author: Tomoki Yoneyama, Address: 350 Massachusetts Ave, Cambridge, MA 02139, USA. Postal address: 350 Massachusetts Ave, Cambridge, MA 02139, USA. Author Contributions T.Y. wrote the manuscript; T.Y., M.K., K.P., and H.W. designed and performed the research and T.Y., H.W. and A.Z.X.Z. analyzed and interpreted the data.

## Abstract

Bispecific T cell engager (TCE) is an emerging anti-cancer modality which redirects cytotoxic T cells to tumor cells expressing tumor-associated antigen (TAA) thereby forming immune synapses to exerts anti-tumor effects. Considering the protein engineering challenges in designing and optimizing size and pharmacokinetically acceptable TCEs in the context of the complexity of intercellular bridging between T cells and tumor cells, a physiologically relevant and clinically verified computational modeling framework is of crucial importance to guide the process to understand the protein engineering trade offs. In this study, we developed a quantitative, physiologically based computational framework to predict immune synapse formation for a variety of molecular format of TCEs in tumor tissue. Our model incorporated the molecular size dependent biodistribution using the two pore theory, extra-vascularization of T cells and hematologic cancer cells, mechanistic bispecific intercellular binding of TCEs and competitive inhibitory interaction by shed targets. The biodistribution of TCE was verified by positron emission tomography imaging of [^89^Zr]AMG211 (a carcinoembryonic antigen-targeting TCE) in patients. Parameter sensitivity analyses indicated that immune synapse formation was highly sensitive to TAA expression, degree of target shedding and binding selectivity to tumor cell surface TAA over shed target. Interestingly, the model suggested a “sweet spot” for TCE’s CD3 binding affinity which balanced the trapping of TCE in T cell rich organs. The final model simulations indicated that the number of immune synapses is similar (∼50/tumor cell) between two distinct clinical stage B cell maturation antigen (BCMA)-targeting TCEs, PF-06863135 in IgG format and AMG420 in BiTE format, at their respective efficacious dose in multiple myeloma patients, demonstrating the applicability of the developed computational modeling framework to molecular design optimization and clinical benchmarking for TCEs. This framework can be employed to other targets to provide a quantitative means to facilitate the model-informed best in class TCE discovery and development.

**Author summary:** Cytotoxic T cells play a crucial role in eliminating tumor cells. However, tumor cells develop mechanisms to evade from T cell recognition. Bispecific T cell engager (TCE) is designed to overcome this issue with bringing T cells to close proximity of tumor cells through simultaneous bivalent binding to both tumor-associated antigen and T cells. After successful regulatory approval of blinatumomab (anti-CD19 TCE), more than 40 TCEs are currently in clinical development with a variety of molecular size and protein formats. In this study, we developed a quantitative computational modeling framework for molecular design optimization and clinical benchmarking of TCEs. The model accounts for molecular size dependent biodistribution of TCEs to tumor tissue and other organs as well as following bispecific intercellular bridging of T cells and tumor cells. The model simulation highlighted the importance of binding selectivity of TCEs to tumor cell surface target over shed target. The model also demonstrated a good agreement in predicted immune synapse number for two distinct molecular formats of TCEs at their respective clinically efficacious dose levels, highlighting the usefulness of developed computational modeling framework for best in class TCE discovery and development.

## 1 Introduction

The last decade witnessed the remarkable potentials of immunotherapies for cancers, including immune checkpoint inhibitors (1, 2), chimeric antigen receptor engineered T cells (3), oncolytic viruses (4) and bispecific T cell engagers (TCE) (5–10). TCE is an emerging anti-cancer modality which redirects cytotoxic T cells to tumor expressing tumor-associated antigen (TAA) thereby forming an immune synapse to activate and cause proliferation of T cells near the tumor cells. The activated T cells then exert anti-tumor effects through release of perforin for membrane perforation followed by granzyme-mediated apoptotic death of the engaged tumor cells. An important advantage of TCE is that it directly activates T cells through cluster of differentiation 3 (CD3) signaling allowing bypassing of major histocompatibility complex (MHC) restrictions and antigen specificity of the T cell receptors (TCR). After successful regulatory approval of catumaxomab (anti-epithelial cell adhesion molecule (EpCAM) TCE, withdrawn in 2017) and blinatumomab (anti-CD19 TCE), more than 40 TCEs are currently in clinical development and over 100 of them are under preclinical development targeting both hematologic malignancies and solid tumors (7).

Due to the complex bispecific mechanism of action of TCEs as well as a lack of target cross reactivity between preclinical animal models, typically mice, and humans for efficacy, computational models provide a crucial means to better understand the efficacy impact of drug-related parameters such as binding affinity to TAA/CD3 and protein format as well as physiological expression levels of TAA/CD3 in target tissues of interest. A number of quantitative computational modeling frameworks has been developed and published within last a few years to quantitatively and translationally address these challenges. It started with modeling of in vitro experimental results of TCE-induced cytotoxicity of tumor cells and simultaneous T cells activation. The observed data was successfully delineated by a mathematical modeling framework which accounting for bispecific binding to cell surface TAA on tumor cells and CD3 on T cells under a wide range of TCE concentrations as well as variable effector to tumor cell (E:T) ratios (11). Jiang et al further demonstrated that in vitro cytotoxicity results are predictive of clinical efficacy of blinatumomab with considering the difference in concentrations of tumor cells and T cells between in vitro experimental condition and patients in vivo (12). Subsequently, modeling frameworks were developed to characterize in vivo responses of TCEs in monkey as well as human peripheral blood mononuclear cells (PBMC) transferred mouse model (13–15). Recently, more integrated, clinically oriented modeling framework has been published which incorporated biodistribution of TCEs as well as tissue expression gradient of target tumor cells and T cells (16, 17) or further connected with a large cancer-immunity cycle model that is verified to immune check point inhibitors (18).

However, most of the modeling frameworks published thus far are not designed to compare different molecular size and protein format of TCEs, and there is a lack of quantitative verification of tissue distribution of TCEs under variable abundance of T cells in different tissues in patients. Owing to the advance in therapeutic protein engineering, next generation TCE candidates are consisted of a variety of molecular sizes and structural domains. Thus, significant differences are expected in drug dispositions and biodistribution of those TCEs. In addition to a huge difference expected in elimination half-lives (typical blood elimination half-life is 2-3 hours for blinatumomab which uses a bispecific T cell engager (BiTE) format (fusion proteins consisting of two single-chain variable fragments) at molecular weight of 5.4 kDa (19), while it is 2-3 weeks for full length antibody at 150 kDa), biodistribution of therapeutic proteins is also reported to be influenced by molecular size (20). Furthermore, although the influence of shed targets on pharmacokinetics (PK)/pharmacodynamics (PD) of targeting biologics has been well acknowledged, biodistribution of shed targets and their impact on PK/PD relationship of TCEs at a site of action has not been fully characterized.

With observing encouraging translational predictability of physiologically based pharmacokinetic (PBPK) models for biodistribution of protein therapeutics, especially for monoclonal antibodies (mAbs) (21–23), two pore theory based extra-vascularization models have been proposed for biodistribution of different molecular size of proteins to account for more detailed physiological tissue distribution processes (24–26). Two pore theory assumed that the tissue vascular endothelium is “porous” and the pore radius on endothelial cells can be classified into two groups, a large amount of small pores (∼4 nm) and fewer number of large pores (∼22 nm) with fractional hydraulic conductance at 0.958 and 0.042, respectively (21, 27). The biodistribution of proteins can be projected by explicitly estimating convectional and diffusional distribution rate constants between vascular space and interstitial space of tissues through either small or large pores considering presence of a circular isogravimetric flow driven by the osmotic pressure. The mathematical modeling framework has been developed and verified by experimental data in preclinical animals and humans to characterize biodistribution to a variety of tissues in a molecular size dependent manner (24–26).

In this study, we have developed a quantitative, physiologically based computational modeling framework for molecular designing of TCEs for a variety of molecular size and protein formats. Our platform model can characterize 1) molecular size dependent biodistribution of TCEs and shed targets in blood, bone marrow, spleen, lymph node under the two pore biodistribution theory, 2) physiologically based extra-vascularization and biodistribution of T cells and tumor cells and 3) mechanistic bispecific binding of TCEs to TAA and T cells with accounting for competitive inhibitory binding to shed targets. The two pore theory biodistribution model was translationally scaled up from mice to patients, and biodistribution of TCE to bone marrow and spleen in patients was verified by clinical positron emission tomography (PET) imaging of [^89^Zr]AMG211, anti-carcinoembryonic antigen (CEA)-targeting TCE in BiTE format (28). Additional model characterization was performed after calibrating the model for multiple myeloma and B cell maturation antigen (BCMA) as a phenotypical disease and target, respectively, to demonstrate the value of this platform. This disease/target combination was selected due to the large amount of literature data on different protein formats of anti-BCMA TCEs under clinical development (e.g. PF-06863135 in IgG format and AMG420 in BiTE format) as well as the abundance of information on shed BCMA in circulating blood and its inhibitory effect against TCE-induced cytotoxicity (29). The model was calibrated for selective binding property between tumor cell surface BCMA and shed BCMA estimated from in vitro cytotoxicity assay of both PF-06863135 and AMG420 in presence of shed BCMA from published literature (29, 30). The calibrated platform was interrogated for key drug-related and system-related parameters for molecular design optimization of TCEs. Moreover, the platform model was applied to simulate and compare amount of immune synapses between T cells and tumor cells in bone marrow, the tumor site of multiple myeloma patients, after treatment of distinct TCEs, PF-06863135 and AMG420, at their respective clinical efficacious dose level to understand the amount of target engagement required in the clinic to treat multiple myeloma.

## 2 Results

### 2.1 Model based characterization of in vitro cytotoxicity of TCEs in presence of shed target

A bispecific TCE binding model for immune synapse formation was developed and applied to in vitro cytotoxicity assays of PF-06863135 and AMG420 reported in literature (29, 30). The detailed description of the mathematical model can be found in the Materials and Methods Section as well as in the Supporting information. A schematic description of model structure is shown in Fig 1(A). The overlay of model-simulated and experimental data of concentration-cytotoxicity relationship after treatment of PF-0683135 and AMG420 in absence or presence of various concentrations of shed BCMA are shown in Fig 2(A) and (B), respectively. The model parameters obtained from the literature or estimated based on observed data are summarized in Table 1. The developed mechanistic bispecific binding model reasonably and quantitatively characterized the concentration-dependent cytotoxicity of tumor cells for both PF-06863135 and AMG420 under a wide range of TCE concentrations while accounting for concentration dependent inhibitory effect of shed BCMA. With fixed parameters of binding affinities to BCMA and CD3, and their expression levels in in vitro systems, the binding affinity to shed BCMA and cytotoxicity related parameters were estimated. It was necessary to estimate baseline shed BCMA expression for PF-0683135 assay to quantitatively delineate concentration-dependent inhibitory effect of shed BCMA, which suggested multiple myeloma cells used in the literature may have produced a small amount of shed BCMA in in vitro assay system. This estimation was not performed for AMG420 assay because only one concentration level of shed BCMA (and absence of shed BCMA) was tested; therefore, binding affinity to shed BCMA and its baseline expression cannot be separately estimated. Nevertheless, an attempt was made to estimate the binding affinity to shed BCMA by AMG420 by assuming the same baseline level of shed BCMA (0.428 nM) from PF-0683135 assay, resulting in only approximately 10% change from the original parameter estimate, suggesting low level of shed BCMA would not cause substantial change in binding affinity estimation in AMG420 assay. Taken together, our model analysis indicated that AMG420 had better selectivity to BCMA over shed BCMA (0.1 nM to BCMA and 5.36 nM to shed BCMA) compared with PF-0683135 (0.04 nM to BCMA and 0.0262 nM to shed BCMA). While maximum tumor killing rate by immune synapse (kkillmax) was very close between PF-0683135 and AMG420 (2.71 and 2.93 1/nM/day), immune synapse concentrations at half maximum tumor killing rate (kkill50) were found to be very different (11.4 and 0.0236 nM) possibly due to inter-laboratory experimental differences.

**Fig 1.**
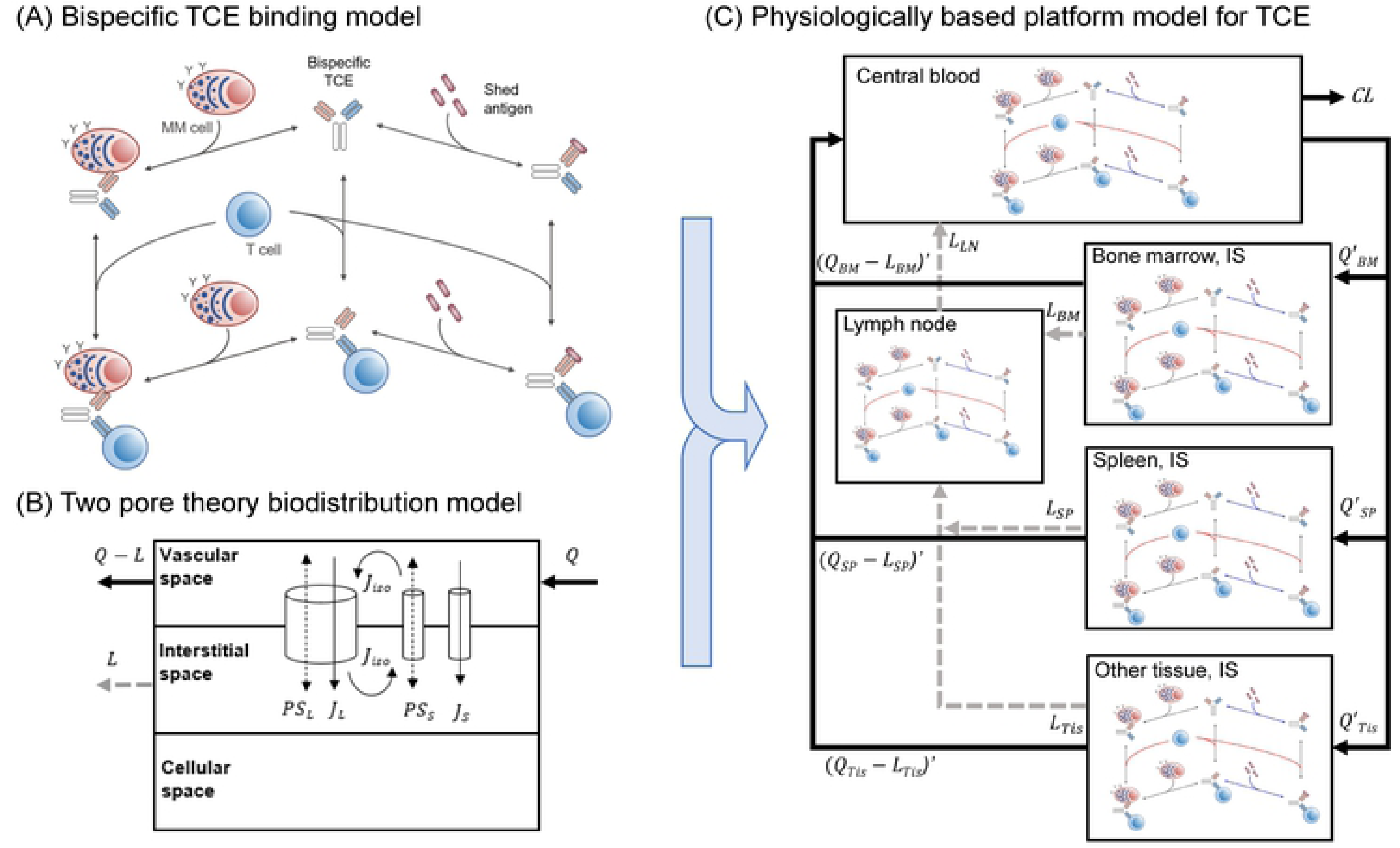
Schematic computational platform model structures for TCE. (A) Mechanistic bispecific binding model of TCE where bispecific TCE binds to either tumor associated antigen (TAA) on tumor cells or CD3 on T cells to form TCE-TAA or TCE-CD3 dimers, followed by subsequent binding with another binding target, either CD3 or TAA, respectively, to form immune synapse (TAA-TCE-CD3 trimer). Shed target competitively inhibits TCE or TCE-CD3 binding to TAA. (B) Two-pore theory biodistribution model. TCE and shed target are transported from vascular space to tissue interstitial space via two groups of pores (small pores: ∼4.44 nm, large pores: ∼22.9 nm) by diffusion and convection in molecular size dependent manner. Q, Q-L and L represents arterial blood flow, venous blood flow and lymphatic flow in each tissue. PS, J and J_iso_ represent diffusion, convection and insogravimetric flow, respectively. Subscript L and S represent large and small pore. (C) Integrated physiologically based platform model for TCE. The biodistribution processes of TCE and shed target into and out of each tissue were characterized by integrated rate constants accounting for perfusion, diffusion, convection and lymphatic flow rate constants in molecular size dependent manner based on two pore theory. Extra-vascularization and biodistribution of T cells and multiple myeloma cells are incorporated into the model. In each tissue and blood compartment, bispecific TCE binding interactions between TAA, CD3 and shed target were considered. Q’ and (Q-L)’ represent integrated biodistribution rate constants in and out of tissue interstitial spaces. L and CL represent lymphatic flow and systemic clearance of TCE or shed target. Subscript BM, SP, Tis, LN represent bone marrow, spleen, lymph node and other tissue, respectively.

**Fig 2.**
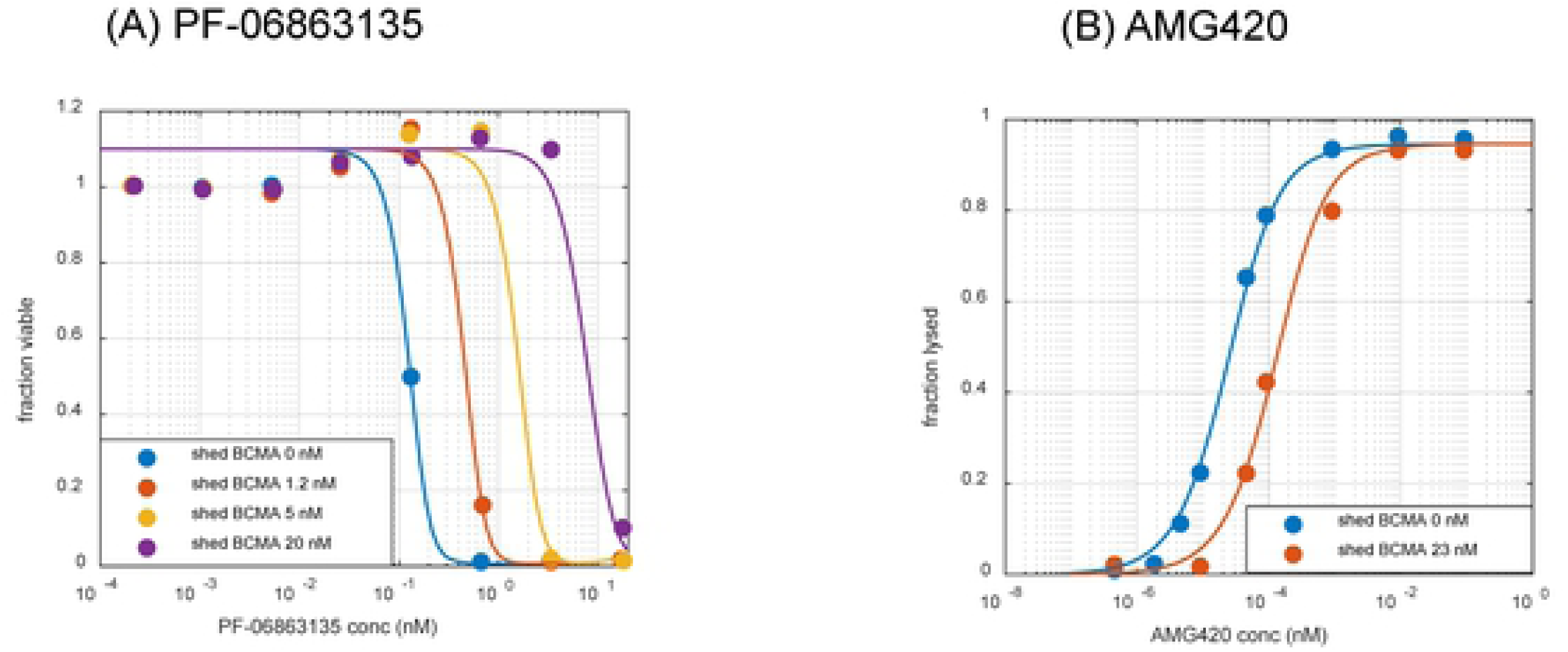
Bispecific TCE binding model-based characterization of in vitro cytotoxicity assay of TCEs. Overlay of experimental observations and model simulation of the relationship between concentration and cytotoxicity for (A) PF-06863135 (IgG format, 150 kDa) and (B) AMG420 (BiTE format, 54 kDa) in absence or presence of various concentration of shed BCMA. In each panel, symbols and lines represent mean observed data and model predictions, respectively. The experimental data were from Panowski et al (29) and Hipp et al (30). sBCMA: shed BCMA.

**Table 1.** Summary of parameters for in vitro cytotoxicity assay for PF-06863135 and AMG420

| Parameter | Unit | Description | Value (%RSE) |  | Source |
| --- | --- | --- | --- | --- | --- |
|  |  |  | PF-06863135 | AMG420 |  |
| kon | 1/nM/day | association rate constant to target, CD3 and shed target | 86 |  | (37) |
| koff | 1/day | dissociation rate constant to target, CD3 and shed target | calc from kon and kd |  |  |
| kd, target | nM | equilibrium dissociation constant to target | 0.04 | 0.1 | *PF-06863135 (29) |
| kd, CD3 | nM | equilibrium dissociation constant to CD3 | 17 | 20 | *AMG420 (42) |
| kd, shed target | nM | dissociation rate constant to shed target | 0.0262 (109) | 5.36 (14.2) | estimated from in vitro data |
| CD3 exp | nM | baseline CD3 expression level | 0.0415 | 0.0498 | PF-06863135, E:T=5:1 (29) |
| BCMA exp | nM | baseline BCMA expression level | 0.00105 | 0.00105 | AMG420, E:T=6:1 (30) |
| sBCMA exp | nM | baseline shed BCMA expression level before addition | 0.428 (23.2) | - |  |
| kkillmax | 1/nM/day | Maximum tumor killing rate by immune synapse | 2.71 (122) | 2.93 (6.7) |  |
| kkill50 | nM | Immune synapse concentration at half maximum tumor killing rate | 11.4 (117) | 0.0236 (12.5) |  |
| hill | - | hill coefficient for tumor killing | 3.09 (32.1) | 1 |  |
| Baseline fraction | - | baseline tumor fraction | 1.10 (1.7) | 1 |  |
| proportional error |  |  | 0.105 | 3E-08 |  |
| additive error |  |  | 0.00854 | 0.0251 |  |

222 \*source for both kd,target and kd,CD

### 2.2 Establishment of molecular size dependent two pore biodistribution model for different size of TCE proteins in mice

The molecular size dependent physiologically based two pore theory model was adapted from literature (24–26) then simplified in following steps to apply to the TCE model: 1) tissues except for bone marrow, spleen and lymph node were lumped into one other tissue, 2) endosomal compartments were removed, 3) tissue vascular spaces were lumped into central blood compartment assuming quasi-equilibrium and 4) biodistribution processes between central blood and tissue interstitial space were simplified under quasi-steady state assumption. More detailed descriptions are found in Materials and Methods as well as Supporting information. Schematic model structure of two pore theory biodistribution model is depicted in Fig 1(B). Schematic model simplification flow of two pore theory biodistribution model is depicted in S1 Fig. The model parameters used and estimated are summarized in S1 and S2 Tables. The overlay of experimental and simplified model simulated concentration-time profiles of domain antibody (dAb_2_, 25.6 kDa) and monoclonal antibody (mAb, 150 kDa) in blood, bone marrow and spleen in mice are described in Fig 3. Without considering endosomal degradation and FcRn-mediated recycling in endosomal compartment, which were incorporated in the original model, the simplified two pore model reasonably delineated the experimental biodistribution data of very different size of proteins in mouse bone marrow and spleen. When impact of tissue lymphatic flow rates was compared using values reported by Sepp et al (25) and Shah et al (23), the former captured experimental biodistribution data better especially for domain antibody in bone marrow and spleen since Shah et al assigned 0.2% of plasma flow rate as lymphatic flow rate, while Sepp et al directly estimated lymphatic flow rate based on tissue distribution of dAb_2_ in mice.

**Fig 3.**
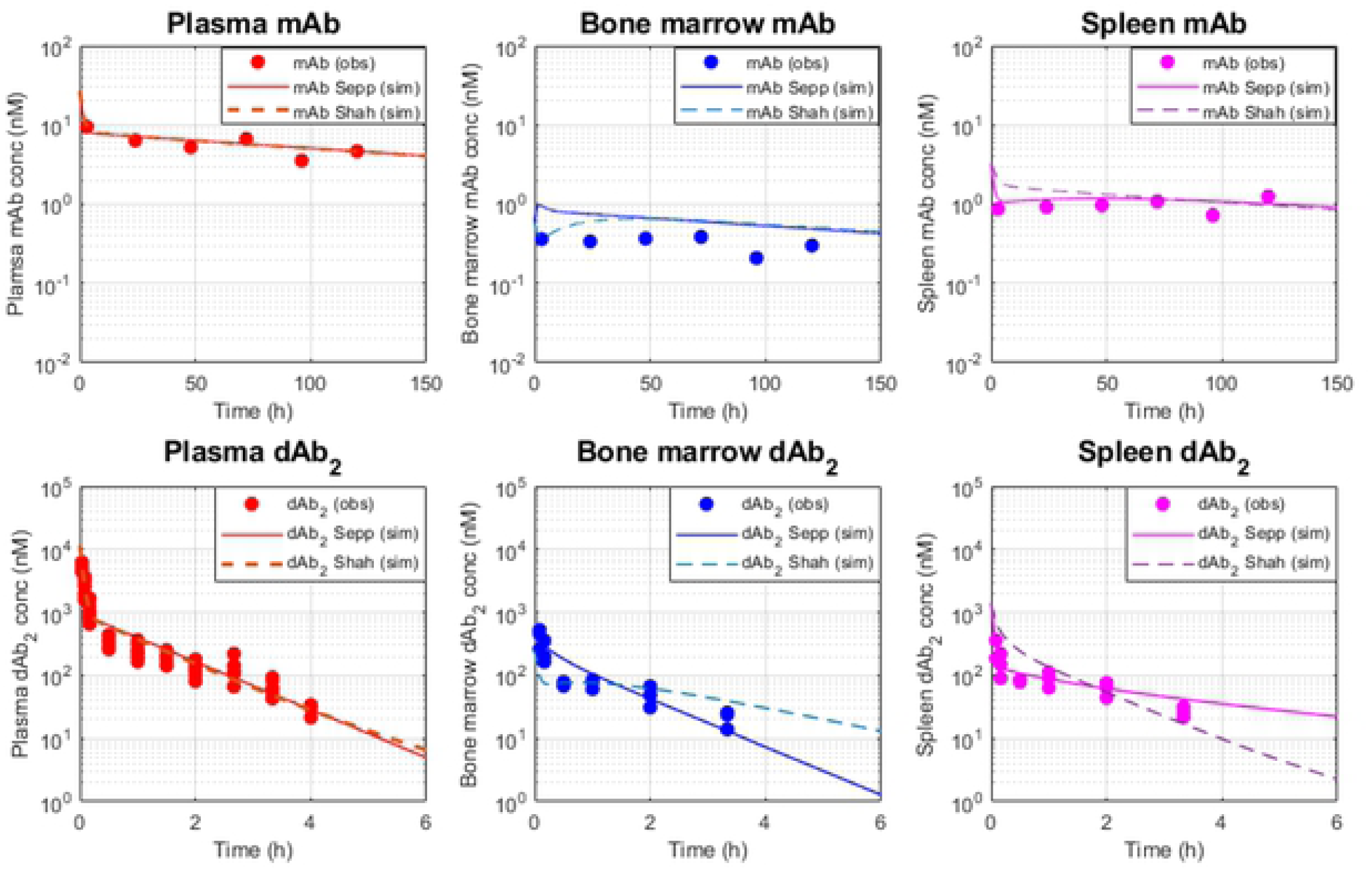
Simplified two pore theory model-based characterization of biodistribution of different molecular size of antibody fragments in mice. Overlay of experimental and model simulation of concentration-time profiles of (A) mAb (150 kDa) and (B) domain antibody (dAb_2_, 25.6 kDa) in plasma, bone marrow and spleen in mice after a single intravenous administration of mAb at 3.8 ug or dAb_2_ at 10 mg/kg in mice. In each panel, symbols represent experimental data. Solid and dash lines represent model predictions under lymphatic flow rate reported by Sepp et al (25) and Shah et al (23), respectively. The experimental data were from Shah et al (23) and Sepp et al (25).

### 2.3 Translation and verification of molecular size dependent two pore theory biodistribution model for TCEs in patients

The simplified two pore model was translated from mice to humans by physiologically scaling up of plasma and lymphatic flow rates as well as tissue volumes. To characterize bispecific TCE binding dispositions, the model was further extended to incorporate T cells extra-vascularization model characterized by Khot et al (31). The developed model was verified using clinical PET imaging data of AMG211, CEA targeting TCE in BiTE format (54 kDa), where biodistribution of AMG211 to tissues including bone marrow and spleen was evaluated (28). The model parameters are summarized in Tables 2 and 3. The overlay of experimental and translated model-simulated concentration-time profiles of AMG211 in blood, bone marrow and spleen in patients are shown in Fig 4. With only estimating systemic clearance (CL) of AMG211, the biodistribution of AMG211 into bone marrow and spleen was reasonably characterized, suggesting that the developed physiologically based two pore theory biodistribution model can be translationally applicable to characterize the biodistribution of different molecular size of TCEs followed by bispecific TCE binding in tissues of interest.

**Fig 4.**
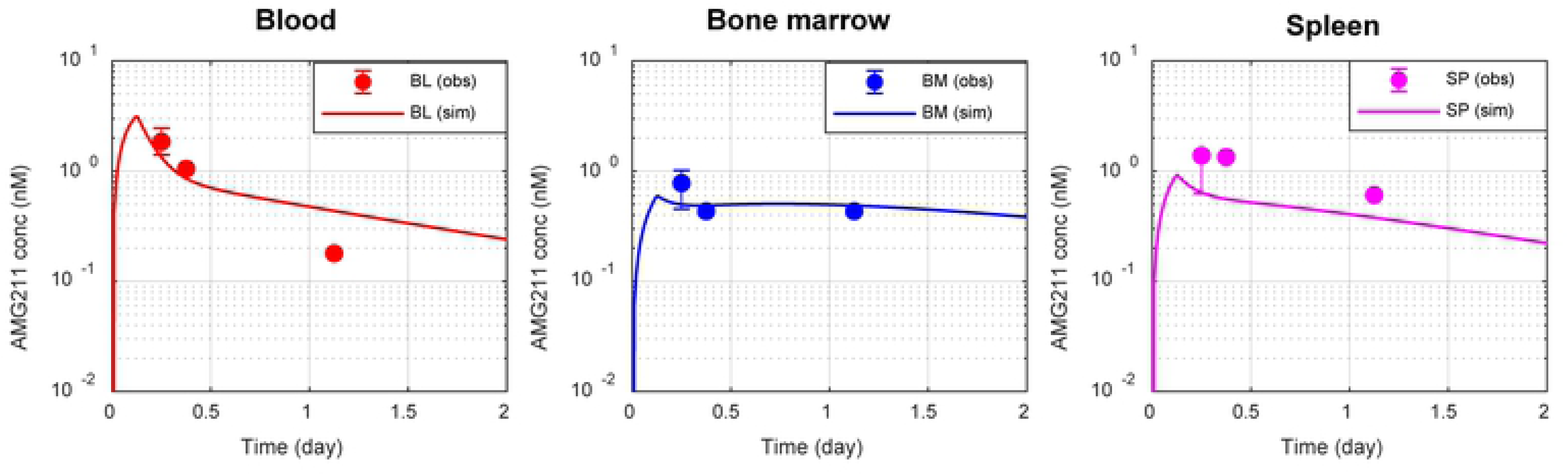
Translated two pore theory model-based characterization of biodistribution of TCE in patients. Overlay of experimental positron emission tomography imaging data and model simulation of concentration-time profiles of AMG211 (carcinoembryonic antigen targeting TCE, BiTE format, 54 kDa) in blood, bone marrow and spleen in patients after intravenous infusion of [^89^Zr]AMG211 at 37 MBq/200 μg with cold AMG211 at 1800 μg for 3 hours. Translated two pore theory model was integrated with T cell extra-vascularization and biodistribution component followed by mechanistic binding of TCE to T cells. Symbols are observed mean for blood and median for tissues (n=4). Lines represent model predictions. The experimental data were from Moek et al (28).

**Table 2.**
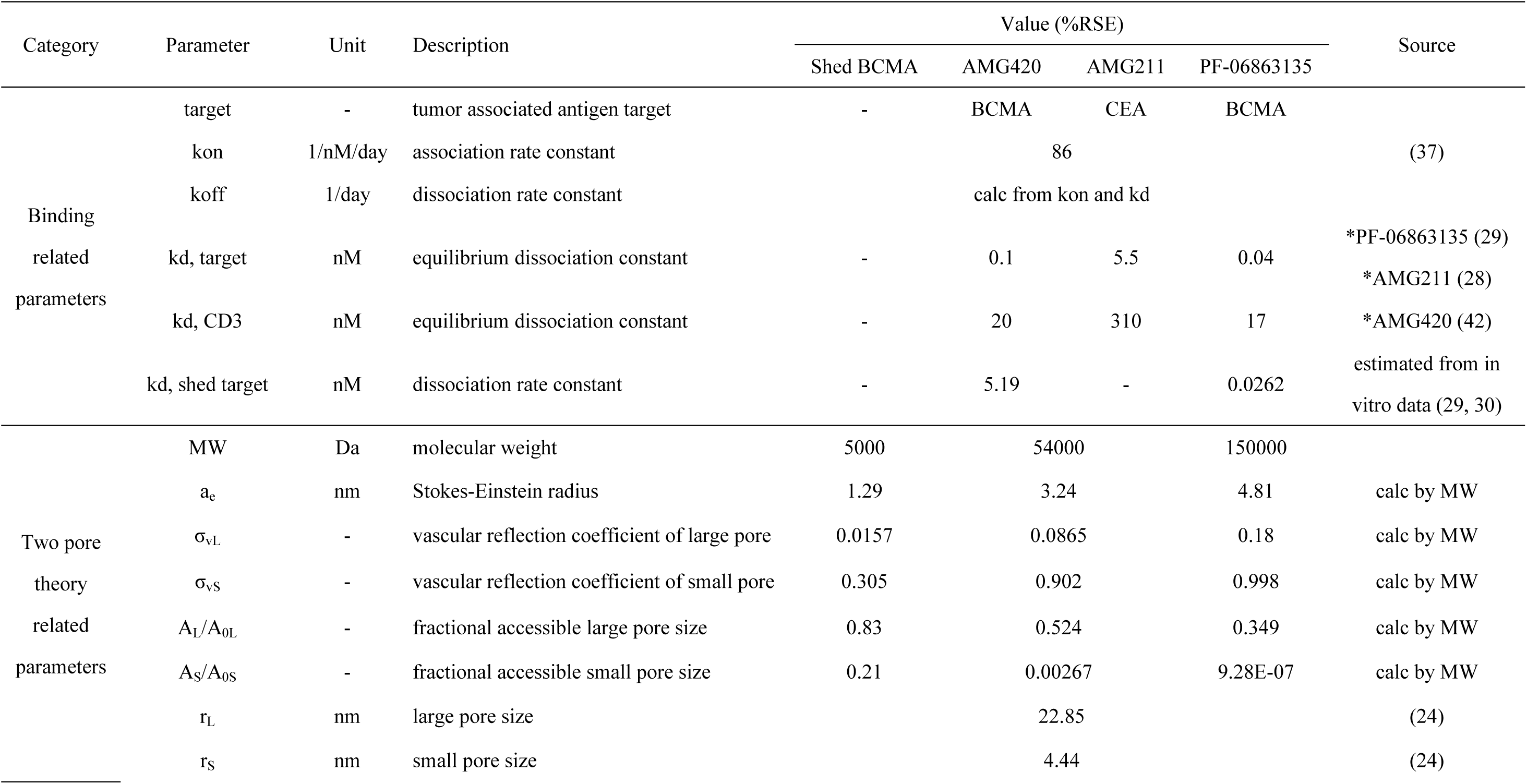
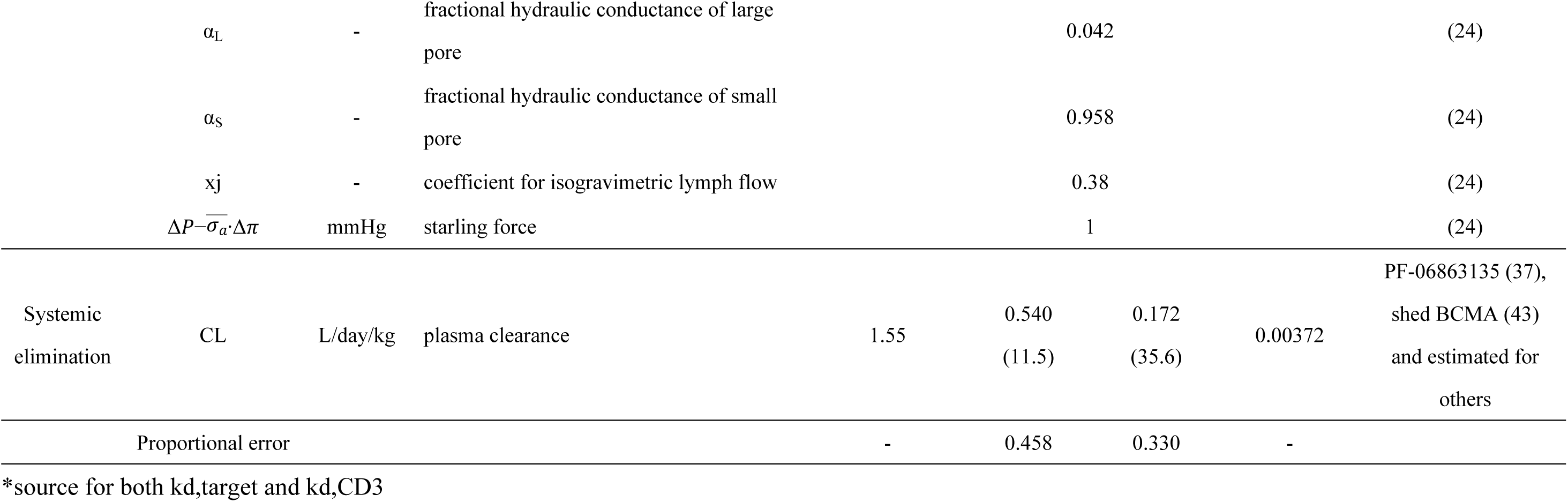
Summary of parameters related to bispecific binding, two pore theory and systemic elimination of shed BCMA, AMG420, AMG211 and PF-06863135 in patients

**Table 3.** Summary of physiological parameters in patients

| Tissue | Total<br>volume<br>(L/kg) | Interstitial<br>volume<br>(L/kg) | Plasma<br>flow<br>(L/day/kg) | Lymphatic<br>flow<br>(L/day/kg) | Lymphatic<br>reflection<br>coefficient | T cell<br>transmigrati<br>on rate<br>(L/day/kg) | BCMA |  | CD3 |  | T cell<br>degradation<br>rate (1/day) | shed BCMA<br>(nM) |
| --- | --- | --- | --- | --- | --- | --- | --- | --- | --- | --- | --- | --- |
|  |  |  |  |  |  |  | conc (nM) | expression<br>(#/cell) | conc (nM) | expression<br>(#/cell) |  |  |
| plasma | 0.044 | - | 61.5 | - | - | - | 4.87E-05 |  | 0.393 |  |  | 100 |
| bone marrow | 0.143 | 0.0266 | 0.88 | 0.0155 | 0.2 | 11.4 | 8.30E-01 |  | 3.66 |  |  | calc by model |
| spleen | 0.0031 | 0.00062 | 2.14 | 0.00018 | 0.2 | 1.88 | 3.49E-02 | 12590 | 98.26 | 100000 | 0.03 | calc by model |
| other tissue | 0.818 | 0.133 | 58.5 | 0.240 | 0.2 | 89.2 | 3.82E-04 |  | 2.21 |  |  | calc by model |
| lymph node | 0.00386 | - | - | 0.256 | - | - | 1.47E-02 |  | 64.5 |  |  | calc by model |
| Source | (23) | (23) | (23) | calc by (23, 25) | (23) | (31) | calc by<br>(32-34) | (35) | calc by<br>(32, 44-46) | (11) | (47) | (38) |

### 2.4 Application of translational platform model to BCMA targeting TCEs in multiple myeloma patients

The verified translated model was calibrated for clinical stage BCMA targeting TCEs, PF-06863135 in IgG format at molecular weight of 150 kDa and AMG211 in BiTE format at molecular weight of 54 kDa. Schematic model structure of physiologically based platform model for TCE is shown in Fig 1(C). The model parameters used and estimated are summarized in Tables 2 and 3. Concentration of BCMA, which is expressed on multiple myeloma cells in blood and different tissues are calibrated based on literature information (32–35). Systemic clearance (CL) of AMG420 was estimated against published clinical PK data (36), while general antibody CL was employed for PF-0683135 (37). The overlay of observed and model simulated plasma concentration time profiles of AMG420 was shown in S2 Fig. The model predicted time profiles of each species (free TCE, free TAA, free CD3, free shed target, dimers and trimers) after treatment of PF-06863135 and AMG420 were shown in Fig 5 and S3 Fig, respectively. The simulation results indicated a dose dependent monotonic increase of free TCE and dimer species (TAA-TCE, CD3-TCE, shed target-TCE). On the other hand, non-monotonic dose response was predicted for immune synapse formation, where dose dependent increase was observed up to approximately 1 mg/kg/week of PF-06863135, while immune synapse levels at 10 and 100 mg/kg/week were lower than 1 mg/kg/week at steady state. Similar non-monotonic relationship in dose to immune synapse formation was predicted for AMG420 as well (S3 Fig). These predicted non-monotonic dose responses in immune synapse formation were consistent with two previous publications on bispecific TCE modeling (14, 18).

**Fig 5.**
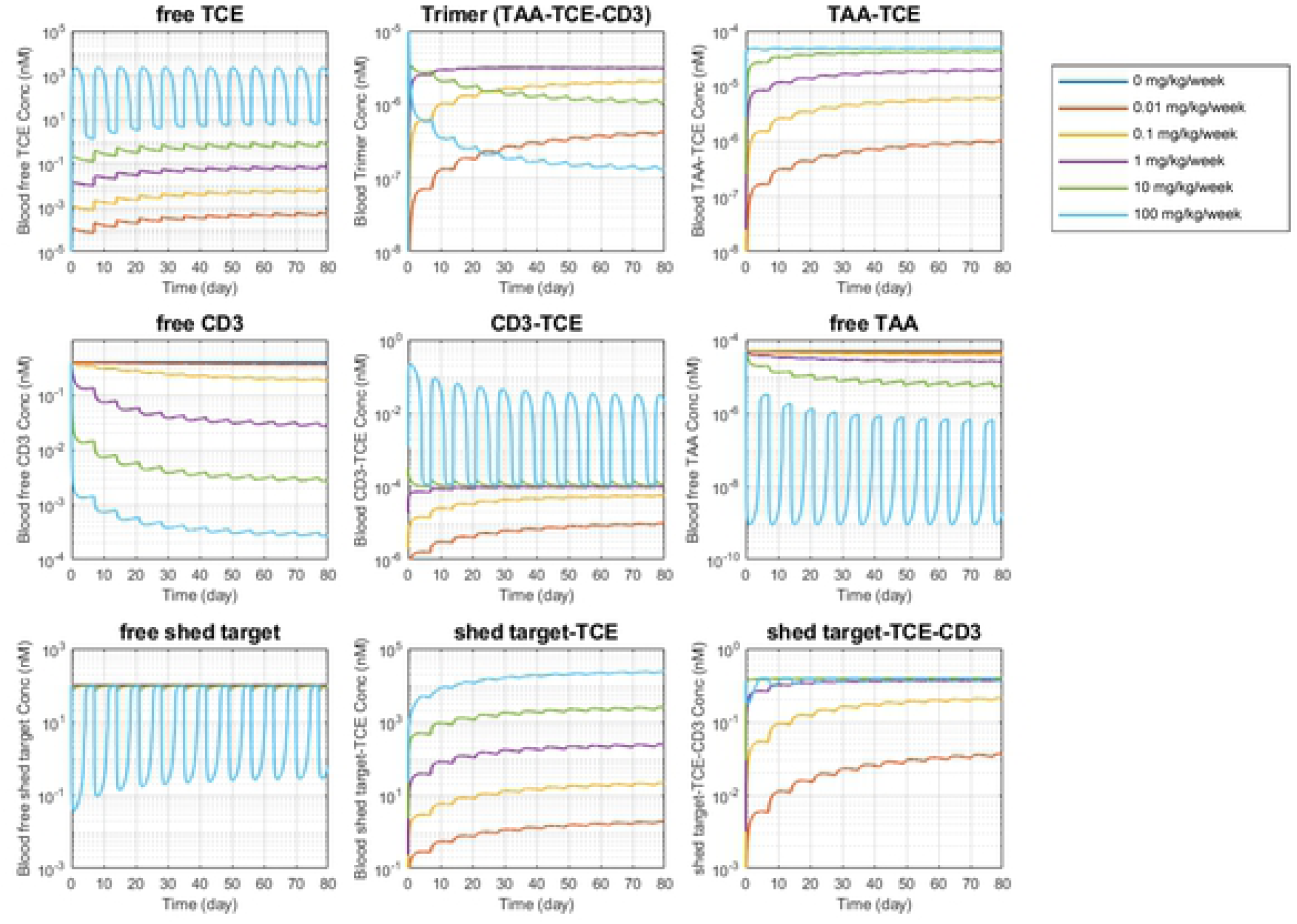
Platform TCE model simulation of concentration time profiles of variable species relating to bispecific TCE binding in blood after subcutaneous administration of PF-06863135 (IgG format, 150 kDa) at 0, 0.01, 0.1, 1, 10 and 100 mg/kg with once weekly dose schedule in multiple myeloma patients. Simulation results were plotted for free TCE, TAA-TCE-CD3 trimer (immune synapse), TAA-TCE dimer, free CD3, TCE-CD3 dimer, free TAA, free shed target, shed target-TCE dimer and shed target-TCE-CD3 trimer in blood.

The developed model of TCEs was further explored using sensitivity analysis to potentially aid protein design of TCEs. The results of local sensitivity analyses were shown in S4 and S5 Figs. Parameters to which immune synapse formation in bone marrow is sensitive for both PF-06863135 and AMG420 with positive correlation were baseline expression of TAA and CD3 in bone marrow, equilibrium dissociation rate coefficient (kd) to shed target. Parameters to which immune synapse formation in bone marrow is sensitive with negative correlation were baseline expression of shed target, kd to TAA and CD3 and elimination rate of shed target. Representing faster elimination of AMG420 due to a lack of FcRn binding domain, systemic clearance was found to be a negatively sensitive parameter only to AMG420. Further sensitivity analysis was performed with wider range of parameter space for key parameters identified from local sensitivity analysis: expression level of and kd to TAA, CD3 and shed target. The results were shown in Fig 6 as well as S6 and S7 Figs. Monotonic relationships were confirmed for expression level of and kd to TAA, CD3 and shed target, except for kd to CD3. These results highlighted the importance of inhibitory relationship between TAA and shed target, suggesting the selectivity between TAA and shed target is one of key factors to maximize immune synapse formation, which enhances anti-tumor effect. On the other hand, intriguingly, a theoretical sweet spot was implied for binding affinity to CD3 (Fig 6). In blood, for example, predicted immune synapse level was monotonically increased with decreasing the kd to CD3 down around 1-10 nM, while immune synapse level started to decrease when further lowering the kd below 0.01 nM.

**Fig 6.**
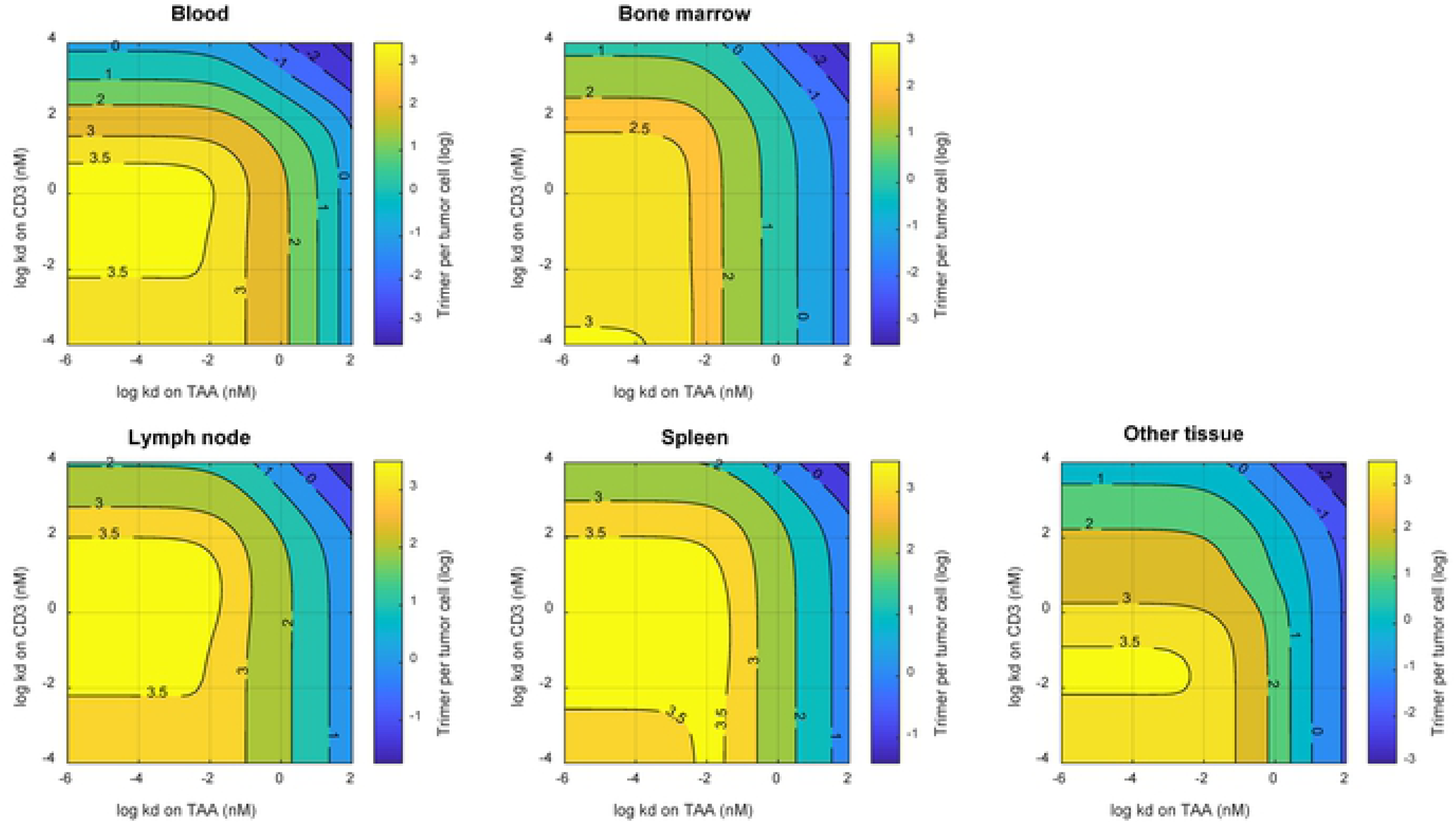
Platform TCE model simulation of two-dimensional contour plots for the effect of TCE’s equilibrium dissociation constants (kd) on TAA and CD3 on immune synapse (TAA-TCE-CD3 trimer) formation in blood, bone marrow, lymph node, spleen and other tissue compartment after subcutaneous TCE (IgG format, 150 kDa) administration once weekly in patients. Other parameters were as with those for PF-06863135.

The platform model simulated dose to immune synapse formation relationships of different molecular size of TCEs, PF-06863135 and AMG420, in absence or presence of shed BCMA were depicted in Fig 7. In absence of shed BCMA, both PF-06863135 and AMG420 reached similar level of maximum immune synapse formation in all the tissue predicted, while PF-06863135 achieved it at lower weekly doses, representing its longer systemic half-life of full IgG format. In presence of shed BCMA (100 nM) at average concentration in multiple myeloma patients (38), meanwhile, maximum level of immune synapse formation was greatly attenuated for PF-0683135, reflecting narrower selectivity window between tumor cell surface BCMA and shed BCMA estimated from in vitro cytotoxicity assay. This prediction emphasizes the importance of selectivity of TCEs to TAA over shed target, which is consistent with the finding from the sensitivity analysis. At the reported clinically efficacious dose of AMG420 (400 ug/day or 0.04 mg/kg/week for 28 days continuous infusion) (36), immune synapse level was predicted to be 53.4/tumor cell in bone marrow. Without any further calibration, the corresponding dose level of PF-06863135, where immune synapse level was at 53.4/tumor cell in bone marrow, was estimated to be 0.302 mg/kg/week under once weekly subcutaneous injection. This projection turned out to be in great agreement with the reported clinically efficacious dose of PF-06863135 (0.215 to 1 mg/kg, once weekly subcutaneous injection) (39). The projected immune synapse level in bone marrow ranged 48.4 to 62.7/tumor cell at reported efficacious dose of PF-06863135, indicating that the projected immune synapse levels in bone marrow were similar between PF-06863135 and AMG420 at their reported efficacious dosing regimen in multiple myeloma patients.

**Fig 7.**
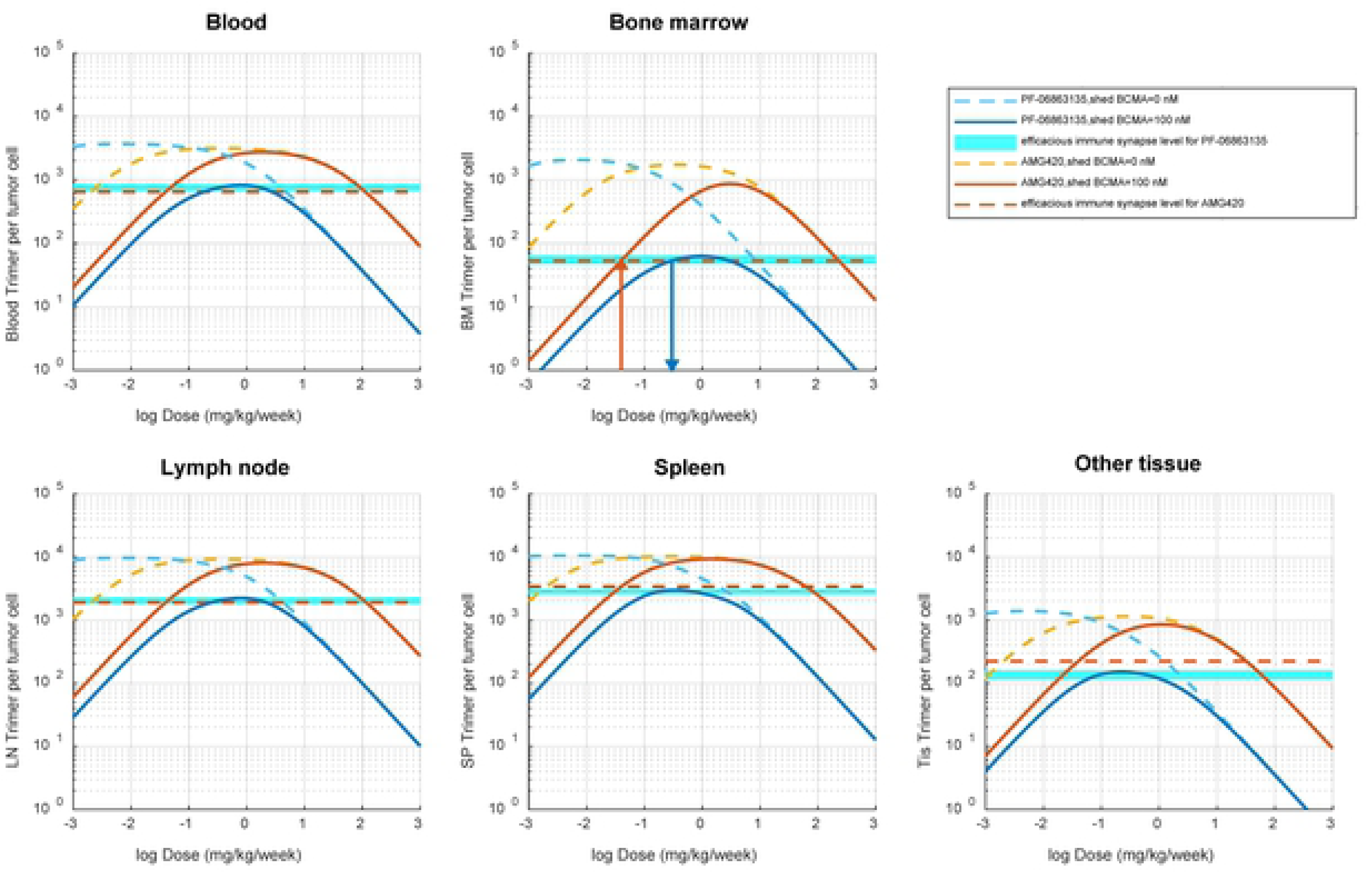
Platform TCE model-based simulation of dose to immune synapse level relationship for PF-06863135 (IgG format, 150 kDa) and AMG420 (BiTE format, 54 kDa) in multiple myeloma patients. Simulation was performed for immune synapse level in blood, bone marrow, lymph node, spleen and other tissue compartment at a wide range of TCE dose levels (0.001 to 1000 mg/kg/week) in absence or presence (100 nM) of shed BCMA in multiple myeloma patients. PF-06863135 was administered once weekly in subcutaneous injections and AMG420 was administered under continuous intravenous infusion for 28 days. Horizontal light blue shade and red dash line represent simulated immune synapse level at reported efficacious dose range of PF-06863135 (250 to 1000 mg/kg/week) and AMG420 (0.04 mg/kg/week), respectively. Red vertical arrow represents predicted immune synapse level in bone marrow at reported efficacious dose level of AMG420 (0.04 mg/kg/week) and bule vertical arrow represents projected dose level of PF-0686135 (0.302 mg/kg/week) at corresponding immune synapse level projected from efficacious dose of AMG420 in patients.

## 3 Discussions

A quantitative computational modeling framework was developed for TCEs with integrating molecular size dependent two pore biodistribution theory, extra-vascularization of T cells and tumor cells as well as mechanistic bispecific binding of TCEs to bridge tumor cells and T cells in presence of shed targets. Although current model consists of blood, bone marrow, lymph node and spleen with reflecting the major tissues of interest for TCE treatment in multiple myeloma patients, the framework can be easily extended to a whole body scale in a full PBPK model. It is also possible to introduce solid tumor compartment by adapting a molecular size dependent tumor disposition model (20). Since there are considerable challenges in applying of TCEs to solid tumor indications, the quantitative modeling framework could potentially provide useful insights for molecular design of TCEs considering unique physio-pathology associated with solid tumor micro-environment as an extension of the present study.

As demonstrated by the modeling results, this modeling framework can be translationally scaled from mice to humans to describe biodistribution of protein therapeutics while considering species specific physiological parameters (e.g. plasma and lymphatic flows, plasma and tissue volumes) as well as abundance of targets, T cells and shed targets. To the best knowledge of the authors, this is the first study to demonstrate that tissue distribution of a TCE in patients could be translationally predicted by two pore theory model combined with T cell abundance in tissues. In light of this finding, the model was further extended for biodistribution of shed targets since competitive interaction of TCEs between membrane surface targets and their shed targets at a site of action is critical to determine the extent of immune synapse formation in the tumor and its downstream pharmacodynamic effect. The developed framework is valuable to predict shed target concentration at a site of action only from circulating level of shed targets and its molecular weight. While there are uncertainties in predicting biodistribution of shed targets, it is important to note that two pore theory biodistribution model has encouragingly broad applicability to not only albumin and antibody fragments but also cytokines (such as IL-2, IL-10, IL-11, etc) and other therapeutic proteins (such as IGF-1, hGH, EPO, etc) (26). Thus, an important future research direction could be charactering the diffusion of protein in tissue interstitial space with respect to its charge, isoelectric point and glycosylation status (24, 26).

The developed modeling framework highlighted the key features of TCE therapeutics that immune synapse formation is a function of binding affinities as well as expression levels of TAA, CD3 and shed targets (Fig 6 as well as S6 and S7 Figs). Especially, the selectivity to tumor cell surface target over shed target is one of the most important molecular designing parameters when the therapeutic target exhibits considerable shedding. As discussed above, the developed platform modeling framework is useful to project competitive interaction of TCEs to target and shed target at a site of action with employing the experimental assay results for binding selectivity estimation.

The developed model implied the existence of a “sweet spot” for binding affinity to CD3, suggesting too tight CD3 binding may not necessarily provide additional benefit on immune synapse formation (Fig 6). The corresponding experimental observation has been reported where higher binding affinity to CD3 negatively affected the systemic exposure of TCEs as it shifted distribution of TCE away from tumor tissue to T cell rich tissues (40). This observation was supported by additional modeling exercise where the implied sweet spot of CD3 binding affinity disappeared when T cell expression level was set uniform across all the tissues and blood (data not shown), suggesting tissue dependent CD3 expression gradient affected the biodistribution and retention of TCEs and immune synapse formation. More importantly, one-compartmental bispecific binding model could not reproduce the non-monotonic effect of CD3 binding affinity on immune synapse formation, suggesting the need of relevant in vivo system or multicompartment mechanistic computational model rather than simple in vitro cytotoxicity assay to aid the optimization of CD3 binding affinity during lead optimization stage of drug development.

The developed model was further applied to understand the amount of immune synapse required to distinct molecular size and formats of TCEs, PF-06863135 in IgG format and AMG420 in BiTE format. The difference in molecular size and protein format seemingly causes substantial difference in PK and tissue distribution disposition. Nevertheless, the final platform model predicted a great agreement in immune synapse level in bone marrow (∼50/tumor cell) at their reported respective efficacious dose level of PF-06863135 (0.215 to 1 mg/kg under once weekly subcutaneous injection (39)) and AMG420 (400 ug/day or 0.04 mg/kg/week under continuous intravenous infusion for 28 days (36)) in multiple myeloma patients (Fig 7). The predicted number of immune synapse was low (∼50/tumor cell) compared with average BCMA expression on multiple myeloma cells (∼12590/tumor cell) (35). This result suggests that model-based simulation of immune synapse level at a site of action can be a good strategy to translationally predict efficacious dose of TCEs in patients. The confidence in the model predictions can be strengthened in case clinical benchmarking insight is available from precedented molecules regardless of molecular size or format since the required information is limited to TCE’s molecular weight, binding affinity to target, CD3 and shed target as well as target and shed target expression levels. Meanwhile, it should be noted that the developed model in present study does not account for dynamics of T cells, tumor cells and shed targets. It has been reported that activated T cells were migrated from blood and proliferated in tissues after treatment of blinatumomab in acute lymphoblastic leukemia (ALL) patients (17). In addition, the depletion of multiple myeloma cells is correspondent with decrease of shed BCMA in multiple myeloma patients (36). These dynamic changes in systems will impact the immune synapse formation and should be carefully incorporated into modeling framework in the future, which will better address possible pathophysiological and pharmacodynamic alterations induced by TCE treatments in patients.

Another important observation from current model is the notably different potency between PF-06863135 and AMG420 observed in vitro cytotoxicity assay despite the good agreement in projected immune synapse formation in bone marrow using reported clinically efficacious doses in patients. The in vitro cytotoxicity related parameters of immune synapse (kkill50) were quite different (11.4 nM for PF-06863135 and 0.0236 nM for AMG420) even when using the same tumor cells, L-363 myeloma cells. The possible reason behind this mismatch might be experimental conditions and sensitivity difference between experiments since these in vitro assay results were collated from different publications likely performed in different laboratories. It has been hypothesized that the source of T cells can generate variability in the results of in vitro cytotoxicity assay of TCEs under co-incubation of tumor cells and effector cells (41). Large sensitivity difference has been also observed between NCI-H929 and L-363 cells when treated with AMG420 in vitro (30). Ideally, in vitro assays performed under controlled setting may provide more quantitative insights. Nevertheless, in vitro assays in presence of shed targets provided useful information to estimate binding affinity/selectivity to shed targets. In addition, Jiang et al reported that in vitro cytotoxicity assays were found predictive for clinical efficacy of blinatumomab (anti-CD19 targeting TCE in BiTE format) in ALL patients (12) with introducing attenuated T cell cytotoxicity (one third of in vitro value) considering immunosuppressive environment in bone marrow in ALL patients compared with in vitro setting. In vitro to in vivo extrapolation of TCE efficacy is obviously the area which needs further investigation. In conclusions, a quantitative computational framework for TCE-mediated immune synapse formation in patients was developed by integrating the two pore theory based biodistribution model of different molecular size of proteins and T cell extra-vascularization model followed by mechanistic bispecific binding in presence of shed target. The developed framework demonstrated the translational predictability of biodistribution of a TCE into bone marrow and spleen in patients, which was supported by PET imaging in clinic. The model framework was particularly useful to explore optimal molecular design parameters of TCEs across a variety of molecular size and formats while considering tissue distribution as well as mechanistic bispecific binding of TCEs to tumor cell surface TAAs and T cells under competitive inhibitory interaction against shed targets. The model highlighted the importance of selectivity to targets on tumor cells over shed targets. The developed modeling framework can quantitatively guide how much selectivity is required in binding affinities in context of expression levels of TAAs and shed targets at a site of action. Moreover, the model implied the existence of sweet spot of CD3 binding affinity, which was not able to be predicted by one compartment model, emphasizing the usefulness of multi-tissue and multi-scale platform models. The model simulated immune synapse levels in bone marrow were found comparable between clinical stage BCMA targeting TCEs, PF-06863135 and AMG420, at reported efficacious dose and dose schedule, suggesting the developed model framework is powerful to leverage clinical benchmarking insights regardless of the difference in molecular size and formats as well as their associated distinct dispositions in pharmacokinetics and tissue distribution. This framework can be applied to other targets beyond multiple myeloma to provide a quantitative means of molecular design and clinical benchmarking in order to facilitate the model-informed best in class TCE discovery and development.

## 4 Materials and methods

### 4.1 Bispecific TCE binding model for in vitro cytotoxicity assay in presence of shed target

Bispecific TCE binding was characterized by a sequential mechanistic binding kinetics model. Schematic description of model structure is shown in Fig 1(A). Differential equations for bispecific binding reactions are shown in Supporting information. The mechanistic binding model was applied to in vitro cytotoxicity assays of PF-06863135 and AMG420 reported in the literature(29, 30). A series of concentrations of PF-06863135 were co-incubated with luciferase-labelled L-363 myeloma cells and human CD3+ T cells at E:T ratio of 5:1 (25000 T cells and 5000 myeloma cells) in absence or presence of shed BCMA at 1.2, 5 and 20 nM. After 2 days of incubation, viability of myeloma cells was assessed by luminescence measurement (29). Likewise, a series of concentrations of AMG420 were co-incubated with L-363 cells and T cells at E:T ratio of 6:1 in absence or presence of shed BCMA at 115 ng/mL (23 nM). The depletion of target cells was measured by flow cytometry analysis after 24 h of incubation (30). Cytotoxicity induced by immune synapse was described by hill function in Eq S10. Parameters used and estimated for in vitro cytotoxicity results were found in Table 1. Target and CD3 expression level were calculated from assay conditions in the literature.

### 4.2 Simplification of molecular size dependent two pore theory biodistribution model for different size of proteins in mice

The biodistribution of different molecular size of TCEs and shed targets was delineated using the two pore theory biodistribution model adapted from the literature (24). Schematic description of two pore theory biodistribution model structure is shown in Fig 1(B). Under two pore theory, proteins are transported from vascular space to interstitial space via two groups of pores (small pores: ∼4.44 nm, large pores: ∼22.9 nm with fractional hydraulic conductance at 0.958 and 0.042, respectively) by diffusion and convection. Equations to determine molecular size dependent transport process under two pore theory can be found in the Supporting information. Original full tissue two pore model was simplified. Detailed model simplification steps are described in Supporting information. Schematic model simplification flow is depicted in S1 Fig. The simplified distribution rate constants in-between central blood and interstitial space (Q’ and (Q-L)’) were derived by integrating perfusion as well as diffusion and convectional rate constants through small and large pores in each compartment as follows;

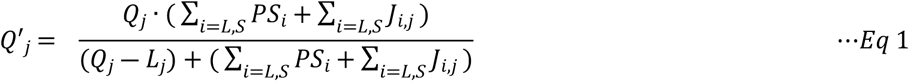

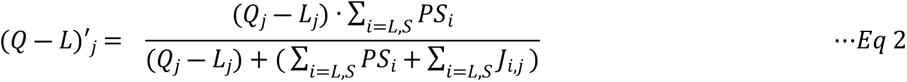

Where Q, (Q-L), PS, J represent arterial plasma flow, venous plasma flow, permeability surface areas, convectional flows, respectively. Subscript L, S and j represent large pore and small pore and jth tissue, respectively.

The simplified two pore biodistribution model was verified using literature information on tissue biodistribution of nonspecific proteins, dAb_2_ (25.6 kDa) and mAb (150 kDa), in mice (23, 25). Parameters used and estimated for in vivo tissue distribution in mice were found in S1 and S2 Tables. Plasma and tissue concentrations of nonspecific mAb and dAb_2_ were evaluated after a single intravenous administration at 3.8 ug and 10 mg/kg, respectively, in mice. Only systemic clearance (CL) was estimated using experimental plasma concentration-time profile data and then simulated tissue concentration profiles using either a set of lymphatic flow rate parameters reported by Sepp et al (25) or Shah et al (23). Shah et al assigned 0.2% of plasma flow rate as lymphatic flow rate, while Sepp et al directly estimated lymphatic flow rate based on tissue distribution of dAb_2_ in mice. The model simulated tissue concentration-time profiles were compared with the experimental results from the literatures.

### 4.3 Translation and verification of two pore biodistribution model for TCEs in patients

The simplified and verified two pore biodistribution model in mice was translationally scaled to humans with replacing the physiological parameters including blood and lymphatic flow rates as well as blood and tissue volumes. The model was further integrated with T cell transmigration/biodistribution model from literature (31) with employing similar model simplification for transmigration of T cells from central blood to tissue interstitial space via tissue vascular space. More details on model translation and T cell transmigration integration processes can be found in Supporting information. Parameters used and estimated for in vitro cytotoxicity results were found in Tables 2 and 3. The translated computational model for bispecific TCE was verified using literature information on biodistribution of AMG211, carcinoembryonic antigen (CEA) targeting TCE, in patients(28). [^89^Zr]AMG211 at 37 MBq/200 μg with cold AMG211 at 1800 μg was intravenously infused to patients for 3 hours. Whole body PET scans were performed at 3, 6 and 24 hours after completion of infusion. Assuming limited CEA expression in blood, bone marrow, spleen and lymph node, kon, target was set to 0 and only CL of AMG211 was estimated. Without further parameter optimization, the biodistributions of AMG211 into bone marrow and spleen were simulated by human model and compared with observed PET imaging data of AMG211 in patients.

### 4.4 Application of translational platform model to BCMA targeting TCEs in multiple myeloma patients

The translated and verified computational model was applied to BCMA targeting bispecific TCEs in multiple myeloma patients. The synthesis, elimination and biodistribution of shed target, shed BCMA, was also incorporated to explicitly account for the competitive interaction against shed BCMA at sites of action. Shed BCMA was assumed to be generated in bone marrow, eliminated from blood circulation mainly through renal excretion and biodistributed to tissues according to two pore theory. Schematic description of final computational platform model structure is shown in Fig 1(C). The final model consists of 45 ordinal differential equations, 197 reactions and 85 parameters. The model parameters used and estimated are tabulated in Tables 2 and 3. Binding affinities of PF-06863135 and AMG420 to shed BCMA were estimated from in vitro cytotoxicity assays and directly employed into the model. Clinical pharmacokinetics of AMG420 was calibrated using literature information on concentration-time profile of AMG420 with 28-day continuous infusion in MM patients. Clearance of PF-06863135 was assumed to be similar to typical antibody (36).

The final platform model was used to simulate concentration-time profiles of variable species relating to bispecific TCE binding in blood, bone marrow, spleen, lymph node and other tissue compartment after administration of PF-06863135 under once weekly subcutaneous injection and AMG420 with continuous intravenous infusion for 28 days in multiple myeloma patients. Local parameter sensitivity analyses were performed around the final sets of model parameters for PF-06863135 and AMG420 to identify parameters to which immune synapse formation is sensitive in blood, bone marrow and lymph node. Additionally, two-dimensional sensitivity analyses were performed to cover wider ranges of key parameters identified from local sensitivity analysis: binding affinity to as well as expression profile of TAA, CD3 and shed target. The relationships between dose of TCEs (0.001 to 1000 mg/kg/week) and immune synapse formation (average from day 56 to 70 for PF-06863135 and average from day 0 to 28 for AMG420) in blood, bone marrow, lymph node, spleen and other tissue compartment were simulated and compared after treatment of PF-06863135 and AMG420 in absence or presence of shed BCMA (100 nM) in multiple myeloma patients.

### 4.5 Model software

The computational model was development using the Simbiology toolbox of MATLAB R2019a (Mathworks, Natick, MA). The optimization toolbox with fminsearch was used with ode15s for parameter estimation. The local parameter sensitivity analyses were performed through calculating time-dependent sensitivity indices on immune synapse formation in blood, bone marrow and lymph node with the full dedimensionalization option and then integrated throughout the time course.

